# On the Robustness of Biomolecular Systems to Perturbations in Translational Resources

**DOI:** 10.64898/2026.08.04.742896

**Authors:** Akshay Kumar Jaiswal, Esha Singh, Abhilash Patel, Soumya Ranjan Sahoo

## Abstract

The reliable operation of biomolecular circuits depends on the availability of shared cellular resources such as ribosomes, whose levels can vary substantially across growth conditions and cellular contexts. Although resource competition among co-expressed genes is well recognized, the relationship between resource variation and the robustness of circuit dynamics has not been characterized quantitatively. This paper integrates a resource-aware gene expression model, contraction theory–based analytical bounds, and experimental validation to study the effect of translational resource variation on constitutive gene expression and its mitigation through feedback. We show that the constitutive circuit exhibits sensitivity to resource perturbations, and that redesigning it with negative autoregulatory feedback enhances the contraction rate and reduces the steady-state deviation bound, though at lower expression levels. Experiments in *E. coli* using both plasmid copy number variation and a ribosome sequestration module are consistent with these predictions, confirming that the feedback circuit maintains relatively stable expression under conditions where the constitutive circuit shows large changes. These findings offer a systematic approach for analyzing and improving the robustness of biomolecular circuits operating under variable resource conditions.

## I. Introduction

Robustness is a critical design requirement in biomolecular systems, as it enables reliable function in the presence of perturbations. It is not a universal property but depends on both the system architecture and the nature of the perturbation encountered [1], [2]. For instance, a circuit that is robust to one class of disturbances may remain sensitive to others [3]. Consequently, understanding and designing robustness to specific perturbations remains a central challenge in synthetic biology [4]. One such perturbation, which is the focus of this work, arises from the limitation and variability of shared translational resources. Within the cell, ribosomes are finite and must be distributed among all expressed genes. This competition gives rise to resource variation, which can alter gene expression levels, may lead to loss of circuit dynamics [5].

Several studies have established that genes compete for a limited pool of ribosomes, creating unintended coupling between circuit components and leading to trade-offs in protein production [6], [7]. The isocost line framework shows that increasing the expression of one gene reduces the resources available to others, constraining overall cellular capacity [7]. Modeling studies have further shown that shared translational resources impose global constraints across genes and significantly affect circuit performance [8], [9]. Resource competition has also been shown to introduce expression burden, reduce modularity, and limit scalability [10]. To address these challenges, approaches such as orthogonal ribosome systems and dynamic resource allocation have been proposed to decouple competing genes and improve predictability [11]. Interestingly, resource limitations can also be actively exploited as a control mechanism to regulate circuit behavior [12]. Additional studies confirm that ribosome loading, cellular burden, and growth-dependent effects significantly influence gene expression and must be accounted for in circuit design [13]–[15].

In this work, we aim to understand how translational resource variation affects the performance of biomolecular circuits and investigate possible design strategies for improving robustness. To address this, we use contraction theory combined with resource-aware gene expression models and experimental measurements. We develop a mathematical model that accounts for ribosomes, a key translational resource, and show that both copy number variations and competition from a sequestering gene result in perturbations in the effective ribosome pool. Using contraction theory, we derive analytical bounds on the trajectory deviation between nominal and resource-perturbed systems and find that the deviation is inversely related to the contraction rate. We analyze the constitutive circuit and identify a fundamental ceiling on its contraction rate imposed by the degradation rate. Next, we show that incorporating negative autoregulatory feedback enhances the contraction rate, resulting in tighter robustness bounds. Finally, experiments are performed in *E. coli* using plasmid copy number variation and a ribosome sequestration module, which show phenomenological agreement with the analytical predictions. These results should help in the design of biomolecular circuits that operate reliably under variable resource conditions.

## II. Mathematical Framework

### A. Preliminaries on Contraction Theory

Contraction theory provides a framework for analyzing how system trajectories evolve relative to one another [16]. In a contracting system, trajectories that start close together converge over time, effectively forgetting their initial conditions. This property makes contraction theory well suited for quantifying robustness: by comparing trajectories of nominal and perturbed systems, one can bound the effect of perturbations on circuit behavior.

Consider a nonlinear system

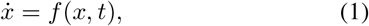

where *x* ∈ ℝ^*n*^ is the state and *f* : ℝ^*n*^ × ℝ_≥0_ → ℝ^*n*^ is a *C*^1^ vector field. The variational (differential) dynamics associated with (1) are

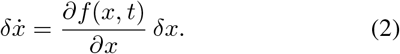

The rate of change of the squared infinitesimal distance between neighboring trajectories satisfies

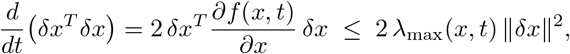

where *λ*_max_(*x, t*) denotes the largest eigenvalue of the symmetric part of the Jacobian, 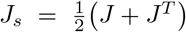, with 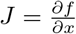.

#### Definition 1.

*[16] The system* (1) *is said to be* contracting *with rate λ >* 0 *if there exists a uniformly positive definite metric M*(*x, t*) = Θ^*T*^ (*x, t*) Θ(*x, t*) *such that the generalized Jacobian satisfies*

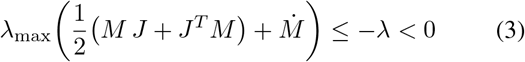

*uniformly for all x and t. In this case*, ∥*δx*∥ _*M*_ *converges to zero exponentially at rate λ, independent of initial conditions*.

#### Remark 1.

*When M* = *I (the identity metric), the contraction condition* (3) *reduces to requiring that the largest eigenvalue of the symmetric part of the Jacobian, λ*_max_(*J*_*s*_), *be uniformly strictly negative*.

### B. Robustness Bounds via Contraction Theory

We now develop bounds on the deviation between trajectories of systems operating under different resource conditions. Consider a biomolecular circuit whose dynamics depend on a resource parameter *r* ∈ ℝ_*>*0_ (e.g., ribosome concentration):

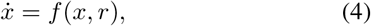

where 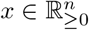 is the state vector and 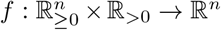 is *C*^1^ in both arguments.

Let two instances of (4) for different resource concentrations of *r*_1_ and *r*_2_:

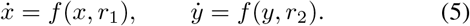

For the deviation *e* ≜ *y* − *x*,

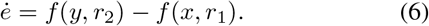

Applying the mean value theorem [17] to the right-hand side, there exist intermediate values 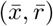 on the line segment joining (*x, r*_1_) and (*y, r*_2_) such that

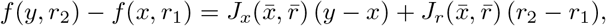

where 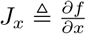 and 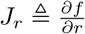. Thus, the deviation dynamics take the form

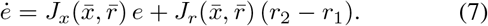

This has the structure of a stable linear system driven by a constant input *r*_2_ − *r*_1_, provided the state Jacobian *J*_*x*_ satisfies a contraction condition. We formalize this as follows.

#### Assumption 1.

*The system* (4) *is uniformly contracting in a metric M* = *M* ^*T*^ *>* 0 *with rate λ >* 0 *over the admissible parameter set* ℛ ⊆ ℝ_*>*0_, *i*.*e*.,

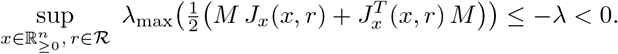

#### Assumption 2.

*The resource sensitivity Jacobian is uniformly bounded: there exists µ >* 0 *such that*

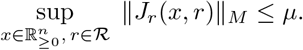

#### Theorem 1.

*Consider the system* (4) *under Assumptions 1 and 2. Let x*(*t*) *and y*(*t*) *be trajectories corresponding to resource parameters r*_1_ *and r*_2_ *with* ∥*r*_2_ − *r*_1_∥ ≤ *δ. Then the deviation e*(*t*) = *y*(*t*) − *x*(*t*) *satisfies:*

i. Transient bound.

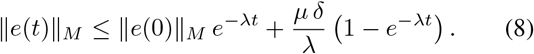
ii. Steady-state bound.

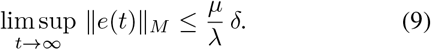

*Proof:* Let ∥ · ∥_*M*_ denote the norm induced by *M*. From the deviation dynamics (7), the contraction condition (Assumption 1) yields the differential inequality

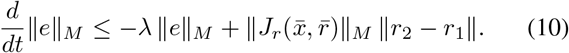

Applying Assumption 2 and ∥*r*_2_−*r*_1_∥ ≤ *δ*, the right-hand side is bounded by −*λ* ∥*e*∥_*M*_ +*µ δ*. By the comparison lemma [17], the solution of the scalar differential inequality 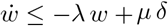 with *w*(0) = ∥*e*(0)∥_*M*_ gives

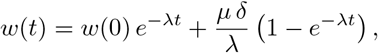

which establishes (8). Taking *t* → ∞ yields (9).

#### Remark 2.

*The bound* (9) *reveals two design levers for improving robustness to resource variation:*

- Increasing the contraction rate *λ. This can be achieved, for instance, by introducing negative autoregulatory feedback, which adds a stabilising term to the Jacobian*.
- Decreasing the resource sensitivity *µ. This corresponds to reducing the dependence of the production kinetics on the resource parameter r*.

#### Remark 3.

*The deviation dynamics* (7) *can be viewed as a stable system with “input”* Δ*r* = *r*_2_ − *r*_1_ *and “state” e. The bound* (9) *is an input-to-state stability (ISS) estimate [18] with gain µ/λ. For a fixed perturbation magnitude δ, a smaller µ/λ implies tighter confinement of trajectories under resource variation. The contraction condition (Assumption 1) guarantees the ISS property, and the steady-state bound is the asymptotic gain*.

### C. Robustness to Resource Variation

We now formalize the notion of robustness to resource variation and introduce quantitative metrics for assessing circuit performance under perturbations.

#### Definition 2

(Robustness to resource variation). *Consider the system* (4) *and let x*(*t*; *r*) *denote the trajectory under resource parameter r* ∈ ℛ ≜ [*r*_min_, *r*_max_].

i. *The system exhibits* perfect resource robustness *over* ℛ *if*

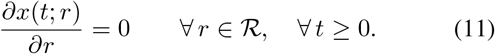
ii. *The system exhibits ϵ*-robustness *over* ℛ *if*

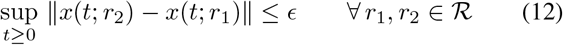

*for a prescribed tolerance ϵ >* 0.

## III. Modeling resource variation co-expression of genes

Consider *i*^*th*^ gene with copy number *n*_*i*_ expresses through the following chemical reactions:

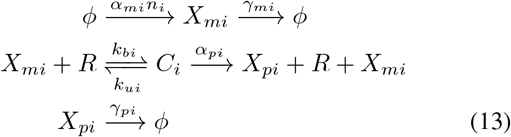

where *X*_*mi*_ is mRNA, *X*_*pi*_ is protein, *R* denotes free ribosomes, and *C*_*i*_ is the ribosome-mRNA complex. Using mass action kinetics, the third-order model can be obtained as,

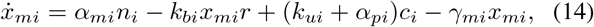

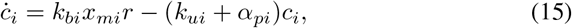

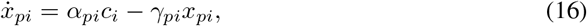

where *x*_*mi*_, *c*_*i*_, *x*_*pi*_, *r* are concentration.

Since ribosome binding/unbinding occurs faster than transcription and degradation, we apply quasi-steady-state (QSS) approximation by setting *ċ*_*i*_ = 0 in Eq. (15),

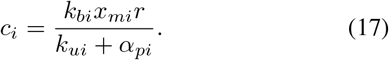

Substituting into Eq. (14), the reduced second-order model becomes:

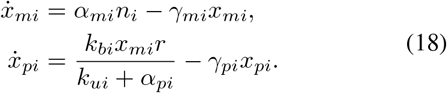

As the total ribosome pool is conserved,

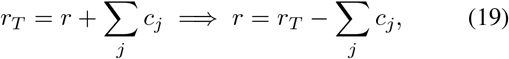

### 1) Resource variation through copy number

For a single gene with copy number *n*, the conservation law (19) reduces to

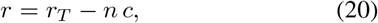

where *c* is the ribosome–mRNA complex concentration per gene copy. Under QSS, 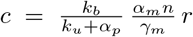, and the free ribosome concentration becomes

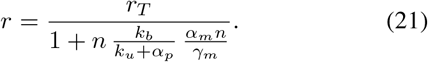

Increasing the copy number *n* raises the total ribosome demand *n c*, reducing the free ribosome pool available for translation.

### 2) Resource variation through a competing gene

Now consider a competing gene with subscript *s* sharing the same ribosome pool. The conservation law becomes

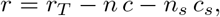

which can be rewritten as

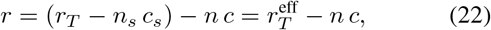

where

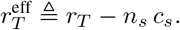

From the perspective of the gene of interest, the competing gene reduces the total available ribosome pool from *r*_*T*_ to an effective pool 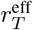. The circuit dynamics retain the same structure as (21) with *r*_*T*_ replaced by 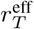:

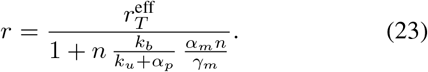

#### Remark 4.

*Both modes of resource perturbation reduce to studying the circuit under a perturbed ribosome pool. Copy number variation changes the demand n c in the denominator, while a competing gene reduces the effective total pool* 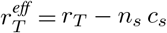. *This equivalence allows both perturbations to be analyzed within the same contraction framework developed in Section II*.

## IV. Parameter Space and Feedback for Robustness

A biomolecular system may show strong robustness for certain parameter values, but become highly sensitive to perturbations when these parameters change. In the previous section, we established that a higher contraction rate implies more robustness.

### A. Constitutive Gene Expression

Consider a mathematical model of gene expression dynamics [12],

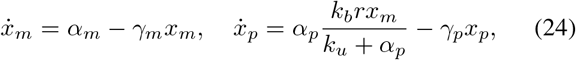

where *x*_*m*_ and *x*_*p*_ are mRNA and protein concentrations, *α*_*m*_ and *α*_*p*_ are production rates, *γ*_*m*_ and *γ*_*p*_ are degradation rates, *k*_*b*_ is the ribosome binding rate, and *k*_*u*_ is the unbinding rate. The symmetrical Jacobian is,

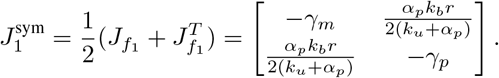

For *γ*_*m*_ = *γ*_*p*_ = *γ*, the eigenvalues satisfy 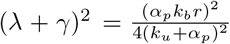, giving the maximum eigenvalue (contraction rate),

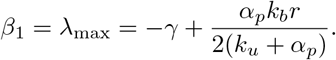

Using the metric of the contraction rate 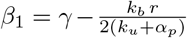, robustness can be improved within the constitutive architecture by increasing the degradation rate *γ*(faster protein turnover), increasing the ribosome unbinding rate *k*_*u*_ (weaker ribosome–mRNA affinity), or decreasing the binding rate *k*_*b*_. However, each of these modifications simultaneously reduces the steady-state protein level, and none can increase the contraction rate beyond *γ* itself. This places a fundamental ceiling on the robustness achievable through parameter tuning alone within the constitutive architecture.

For the constitutive circuit, we numerically evaluated robustness over the parameter plane spanned by *α*_*p*_*/γ* and *k*_*b*_*/k*_*u*_. At each grid point, we fixed a nominal ribosome level *r*_0_, generated 100 random perturbations with uniformly distributed ± 30% variations in *r*, and computed the worst-case value of sup_*t*_ ∥*x*(*t*; *r*_pert_) − *x*(*t*; *r*_0_)∥ . As shown in Fig. 1b, the robustness metric increases monotonically as either *α*_*p*_*/γ* or *k*_*b*_*/k*_*u*_ increases, indicating that the constitutive circuit becomes more sensitive to ribosome fluctuations in these regimes. The most robust region appears in the lower-left corner, where both ratios are small, whereas the upper-right region corresponds to the least robust behavior, reflecting the tradeoff between stronger expression and reduced robustness to resource perturbations.

**Fig. 1.**
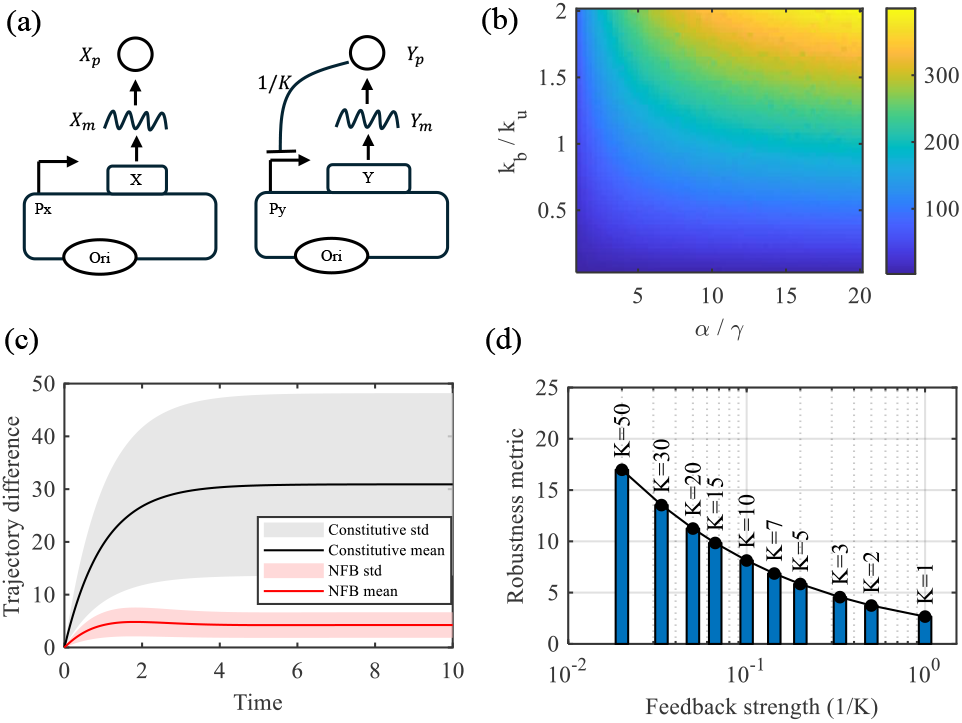
Robustness analysis of constitutive and negative autoregulatory (NAR) circuits under resource variation. (a) Circuit schematics: constitutive expression (top) and NAR with protein-mediated transcriptional repression (bottom). (b) Heatmap of the robustness metric sup_*t*_ ∥*x*(*t*; *r*_pert_) − *x*(*t*; *r*_0_) ∥ for the constitutive circuit over the parameter plane (*α*_*p*_*/γ, k*_*b*_*/k*_*u*_), computed from 100 uniformly distributed ± 30% perturbations in *r*. Lower values (darker) indicate greater robustness. (c) Mean trajectory deviation (solid) and standard deviation band (shaded) for the constitutive and NAR circuits under the same 100 ribosome perturbations. (d) Robustness metric as a function of feedback strength 1*/K* for the NAR circuit, showing that stronger feedback (smaller *K*) improves robustness.

### B. Improving Robustness through Negative Autoregulation

The analysis above shows that the constitutive circuit has a contraction rate bounded above by *γ*, limiting its robustness to resource variation. It has been reported that negative autoregulatory feedback can improve the contraction rate of biomolecular circuits [19]. Motivated by this, we consider redesigning the constitutive circuit by introducing negative autoregulatory feedback, where the protein product represses its own transcription.

The resulting dynamics are

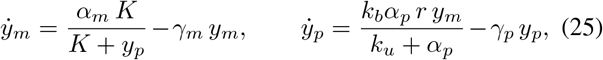

where *K* is the dissociation constant. The transcription rate *α*_*m*_*K/*(*K* + *y*_*p*_) is a decreasing function of the protein level *y*_*p*_, forming a negative feedback loop.

The symmetrical part of the Jacobian is,

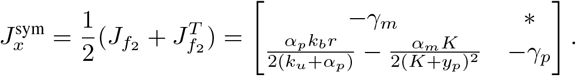

For *γ*_*m*_ = *γ*_*p*_ = *γ*, the maximum eigenvalue is,

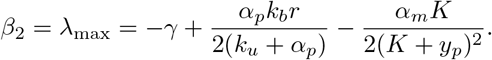

The contraction rate increases by 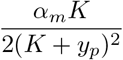 relative to the constitutive circuit, making *β*_2_ more negative. Thus, negative autoregulatory feedback exhibits enhanced robustness to ribosome variation compared to unregulated expression.

To compare robustness under identical resource fluctuations, we generated 100 ribosome perturbations uniformly from ± 30% of the nominal value *r*_0_, and used the same perturbation set for both the constitutive and negative autoregulatory feedback circuits. For each perturbation, we computed the trajectory deviation from the corresponding nominal trajectory, and then plotted the mean and standard deviation across all 100 perturbations. As shown in Fig. 1, the negative autoregulatory feedback circuit exhibits a smaller mean trajectory deviation and a narrower standard-deviation band than the constitutive circuit under the same perturbations. This indicates that negative autoregulation reduces both the average sensitivity and the variability of the response to ribosome fluctuations, consistent with enhanced robustness.

To quantify the effect of feedback strength on robustness, we simulated the negative autoregulatory feedback circuit for a sequence of dissociation constants *K*, using the same 100 ribosome perturbations drawn uniformly from ± 30% of the nominal value *r*_0_ for each case. For every *K*, we computed the robustness metric 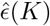 so that smaller values indicate greater robustness. As shown in Fig.1d, the robustness metric decreases as the feedback strength increases, i.e., as *K* decreases (equivalently, as 1*/K* increases). This confirms that stronger negative autoregulation suppresses the effect of ribosome perturbations more effectively, leading to improved robustness.

## V. Experimental Assessment of Resource variation

### A. Materials and Methods

#### 1) Plasmids and Bacterial strains

In this study, negative autoregulatory feedback circut is realised by Ptet/TetR with TetR:GFP fusion protein regulated with Ptet promoter. For constitutive circuit realisation, GFP is regulated with Ptet promoter, but without TetR gene. GFP is used as reporter protein. For ribosome sequestration, mCherry with arabinose was used. These circuits were cloned into p15A vector for low copy and ColE1 vector for high copy implementation. *E. coli* DH5*α* strain was transformed with these circuits.

#### 2) Measurements

The culture was grown overnight by inoculating single colony in LB rich media supplemented with appropriate antibiotic at 37°C, 240 rpm, for 16 hours. Overnight liquid culture was then diluted to 1:50 in fresh M9CA containing appropriate antibiotics. After dilution, culture was induced using given concentrations of inducers (aTc and arabinose) depending on the circuit. Culture was then incubated (37°C, 240 rpm) for 10 hours and point reading was taken at the interval of 2 hours. The sample (200 *µL*) was loaded in 96-well plate in triplicates and point reading was taken using a plate reader where optical density (OD_600*nm*_), GFP Fluorescence (ex./em. = 485/510nm, gain 80), and mCherry Fluorescence (ex./em. = 587/610nm, gain 100) was measured for quantitative analysis. For background subtraction, prior to each reading, blank plate reading was recorded along with absorbance and fluorescence of media alone. These measurements were repeated for three days at 37°C for all experiments.

#### 3) Data Analysis

Data analysis was done using MATLAB. In all plots, bars represent the mean values, and error bars indicate the standard deviation across biological replicates.

### B. Resource variation by changing Copy Number

To assess the impact of resource-demand variation on circuit performance, we compared the expression profiles of constitutive and negative autoregulatory feedback circuits carried on low-copy (p15A) and high-copy (ColE1) plasmids (Fig. 2a). In response to the increase in plasmid copy number, the constitutive circuit exhibited an approximately 20-fold increase in GFP/OD. In contrast, the negative autoregulatory feedback circuit showed only an approximately 1.5-fold change between the two copy-number conditions, indicating greater robustness to plasmid copy-number variation and, consequently, to changes in resource demand.

**Fig. 2.**
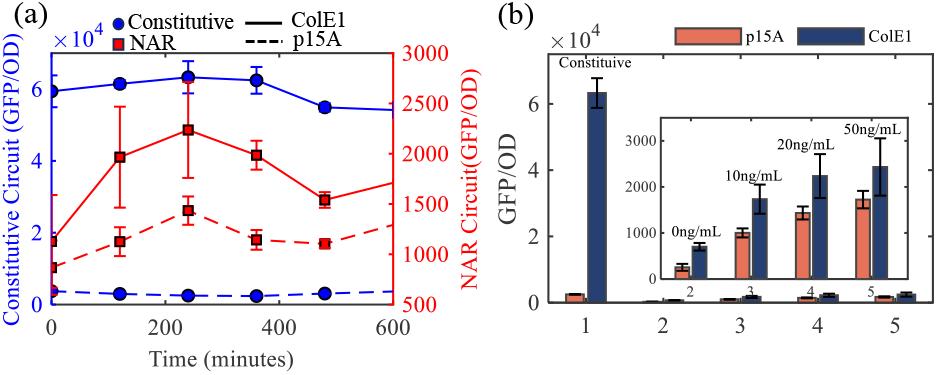
Experimental comparison of constitutive and negative autoregulatory feedback circuits under copy number variation in *E. coli*. (a) GFP/OD over time for constitutive and negative autoregulatory feedback (NAR) circuits on low-copy (p15A) and high-copy (ColE1) plasmids at 20 ng/mL aTc. The constitutive circuit shows an approximately 20-fold increase from p15A to ColE1, while the negative feedback circuit shows only an approximately 1.5-fold change. (b) GFP/OD at *t* = 4 h across aTc concentrations (0, 10, 20, 50 ng/mL) for both copy number conditions.

A similar trend was observed across all tested aTc concentrations (0, 10, 20, and 50 ng/mL), where the constitutive circuit remained highly sensitive to copy number, whereas the negative autoregulatory feedback circuit exhibited only minimal changes in GFP/OD (Fig. 2b). Together, these results suggest that negative autoregulatory feedback improves robustness to resource variation arising from changes in plasmid copy number.

### C. Variation in Resource Availability by Introducing a Sequestration Circuit

To examine the effect of reduced resource availability on circuit performance, a sequestration module was co-transformed with the constitutive and negative autoregulatory feedback circuits (Fig. 3a,b), hereafter referred to as the sequester-constitutive and sequester-negative-feedback circuits, respectively. The sequestration module imposes an additional translational load by drawing from the available ribosome pool, thereby creating a resource-limited intracellular environment.

**Fig. 3.**
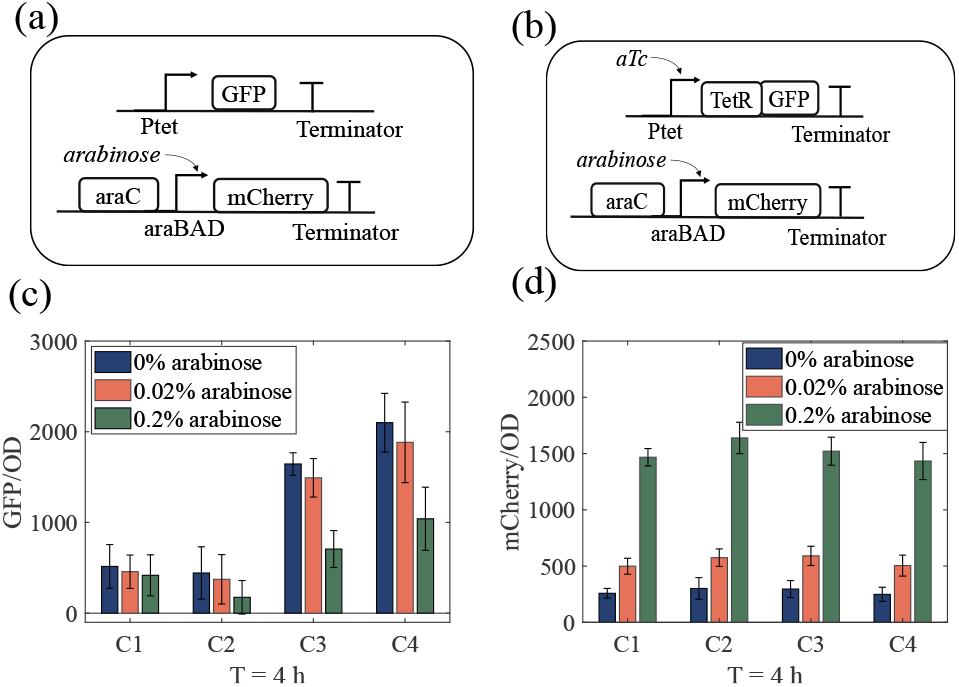
Response of circuits under ribosome sequestration. C1: sequester-constitutive circuit; C2: sequester-negative autoregulatory feedback at 0 ng/mL aTc; C3: sequester-negative autoregulatory feedback at 20 ng/mL aTc; C4: sequester-negative autoregulatory feedback at 50 ng/mL aTc. (a) Schematic of the sequester-constitutive circuit, where an arabinose-inducible mCherry module competes for ribosomes with the constitutive GFP circuit. (b) Schematic of the sequester-negative feedback circuit, where the same sequestration module competes with the TetR-based NAR circuit. (c) GFP/OD at *t* = 4 h under varying arabinose concentrations (0%, 0.02%, 0.2%). Increasing arabinose reduces GFP/OD across all circuits, with the constitutive circuit (C1) showing the largest decrease. (d) mCherry/OD at *t* = 4 h, confirming activation of the sequestration module with increasing arabinose. The negative autoregulatory feedback circuits (C2–C4) maintain higher GFP/OD than the constitutive circuit under comparable sequestration conditions.

To quantify the effect of this imposed load, GFP/OD and mCherry/OD were measured at different time points (0, 4, and 8 h) under varying aTc (0, 20, and 50 ng/mL) and arabinose (0%, 0.02%, and 0.2%) concentrations. At *t* = 4 h, induction of the sequestration module produced a clear inverse relationship between GFP/OD and mCherry/OD across the tested conditions. We noted increasing arabinose concentration increased mCherry/OD (Fig. 3d) while decreasing GFP/OD (Fig. 3c). This trend indicates activation of the sequestration module and the onset of competition for a finite pool of ribosomes.

Notably, the sequester-negative autoregulatory feedback circuit maintained higher GFP/OD than the sequester-constitutive circuit under comparable sequestration conditions, suggesting that negative autoregulatory feedback partially buffers the effect of resource limitation. Overall, these results indicate that negative autoregulatory feedback enhances circuit robustness under reduced resource availability by maintaining more stable gene expression despite sequestration-induced ribosome competition.

## VI. Conclusion

Understanding how biomolecular circuits respond to fluctuations in shared cellular resources is essential for developing and designing predictable synthetic biological systems. We addressed the problem of robustness to translational resource variation in biomolecular circuits through modeling, contraction analysis, and experimental validation. We first modeled ribosome sharing among co-expressed genes under a conservation law and showed that both copy number variation and competition from a second gene reduce to perturbations in the effective ribosome pool. Using contraction theory, we derived analytical bounds on the trajectory deviation between nominal and resource-perturbed systems. For the constitutive circuit, we showed that the contraction rate is bounded above by the degradation rate, placing a ceiling on the robustness achievable through parameter tuning alone. Introducing negative autoregulatory feedback was shown to enhance the contraction rate, yielding tighter robustness bounds at the cost of reduced expression levels. Numerical simulations confirmed that stronger feedback further improves robustness. Experimental measurements in *E. coli* using two independent modes of resource perturbation, plasmid copy number exchange and ribosome sequestration, are consistent with these predictions. The constitutive circuit exhibited large changes in expression under both perturbations, while the negative autoregulatory feedback circuit maintained relatively stable output across all tested conditions. These results offer a systematic approach to analyzing and designing biomolecular circuits that are less sensitive to translational resource variation.

